# High-resolution single-molecule long-fragment rRNA gene amplicon sequencing for uncultured bacterial and fungal communities

**DOI:** 10.1101/2021.03.29.437457

**Authors:** Chao Fang, Xiaohuan Sun, Fei Fan, Xiaowei Zhang, Ou Wang, Haotian Zheng, Zhuobing Peng, Xiaoqing Luo, Ao Chen, Wenwei Zhang, Radoje Drmanac, Brock A. Peters, Zewei Song, Karsten Kristiansen

## Abstract

Although several large-scale environmental microbial projects have been initiated in the past two decades, understanding of the role of complex microbiotas is still constrained by problems of detecting and identifying unknown microorganisms^1-6^.Currently, hypervariable regions of rRNA genes as well as internal transcribed spacer regions are broadly used to identify bacteria and fungi within complex communities^7,8^, but taxonomic and phylogenetic resolution is hampered by insufficient sequencing length^9-11^. Direct sequencing of full length rRNA genes is currently limited by read length using second generation sequencing or sacrificed quality and throughput by using single molecule sequencing. We developed a novel method to sequence and assemble nearly full length rRNA genes using second generation sequencing.Benchmarking was performed on mock bacterial and fungal communities as well as two forest soil samples. The majority of rRNA gene sequences of all species in the mock community samples were successfully recovered with identities above 99.5% compared to the reference sequences. For soil samples we obtained exquisite coverage with identification of a large number of putative new species, as well as high abundance correlation between replicates. This approach provides a cost-effective method for obtaining extensive and accurate information on complex environmental microbial communities.

---

Currently, there are two main approaches for sequencing rRNA genes in complex microbial environmental samples: second generation technologies such as sequencing by synthesis and DNA nanoball sequencing (DNBseq), and third-generation technologies such as nanopore and single molecule real-time sequencing. Second-generation sequencing suffers from relatively short read lengths (less than 300 bases) making it difficult to sequence the entire rRNA gene. Recently, co-barcoding technologies combined with second generation sequencing and *de novo* assembly have enabled nearly full-length rRNA genes, but their quantification reproducibility using complex samples is unknown^9, 10^. Third generation technologies have much longer read lengths enabling full coverage of the rRNA genes, but they are still hampered by relatively high cost, low throughput, and high error rates^11-13^. As such, there is still a great demand for a high-throughput and cost-effective approach to support the deciphering of complex microbial communities. In this study, we developed a strategy whereby a novel combination of rolling circle replication and DNA co-barcoding ^14^ techniques allows sequence information of long amplicons to be obtained by a high-throughput, single-molecule level *de novo* assembly approach to achieve species-resolution and highly accurate profiles in an efficient and cost-effective manner using a short read sequencing platform. To demonstrate the applicability of this method, we benchmarked it using two mock communities of various bacterial and fungal species. We also applied it to field soil samples as a demonstration for discovering unknown species and were able to recover rRNA sequences even longer than those found in current reference databases.

The basic idea is to assemble a single DNA molecule by using DNA co-barcoding technology enabled by the single-tube long fragment read (stLFR) method. Using stLFR, it is possible to label DNA sub-fragments from a single long DNA molecule with the same barcode. Importantly, in a single stLFR library there are approximately 3.6 billion different barcodes allowing each long DNA molecule in a sample to be labeled by a unique barcode^15^. One shortcoming of the current stLFR method is that sequence coverage of each long molecule is less than 1X. As a result, using the De Bruijn graph (DBG) algorithm, it is usually impossible to achieve full assembly of DNA fragments from a single barcode. To overcome this lack of co-barcoded sequence coverage, we applied a modified version of the stLFR technology, named stcLFR (<u>s</u>ingle tube <u>c</u>omplete-coverage <u>L</u>ong <u>F</u>ragment <u>R</u>ead). This approach involves an initial process of rolling-circle replication (RCR), which performs a linear amplification process whereby tandem copies of a single DNA molecule are joined head to tail in one long single stranded concatemer. This method also avoids cumulative replication errors since the same DNA fragment is used as the template for replication. After conversion to double stranded DNA, the RCR amplified products are disrupted at regular intervals of approximately 500 base pairs by a transposon containing a capture sequence. The transposon inserted molecules are then captured by barcode containing oligonucleotides anchored to the surface of a micron sized magnetic bead, whereupon the DNA is fragmented and ligated to these oligonucleotides (Figure 1a). After PCR on the beads, the DNA fragments with unique barcodes, corresponding to multiple copies of each single molecule, are subjected to DNB based sequencing. By decoding barcoded sequences from the sequencing dataset, copy depth can be estimated in each barcoded binning group. Bins with depth > 5X are then used for parallel *de novo* assembly (Figure 1b). Successfully assembled contigs are termed pre-binning assemblies (PBAs) and used for subsequent analysis.

**Figure 1.**
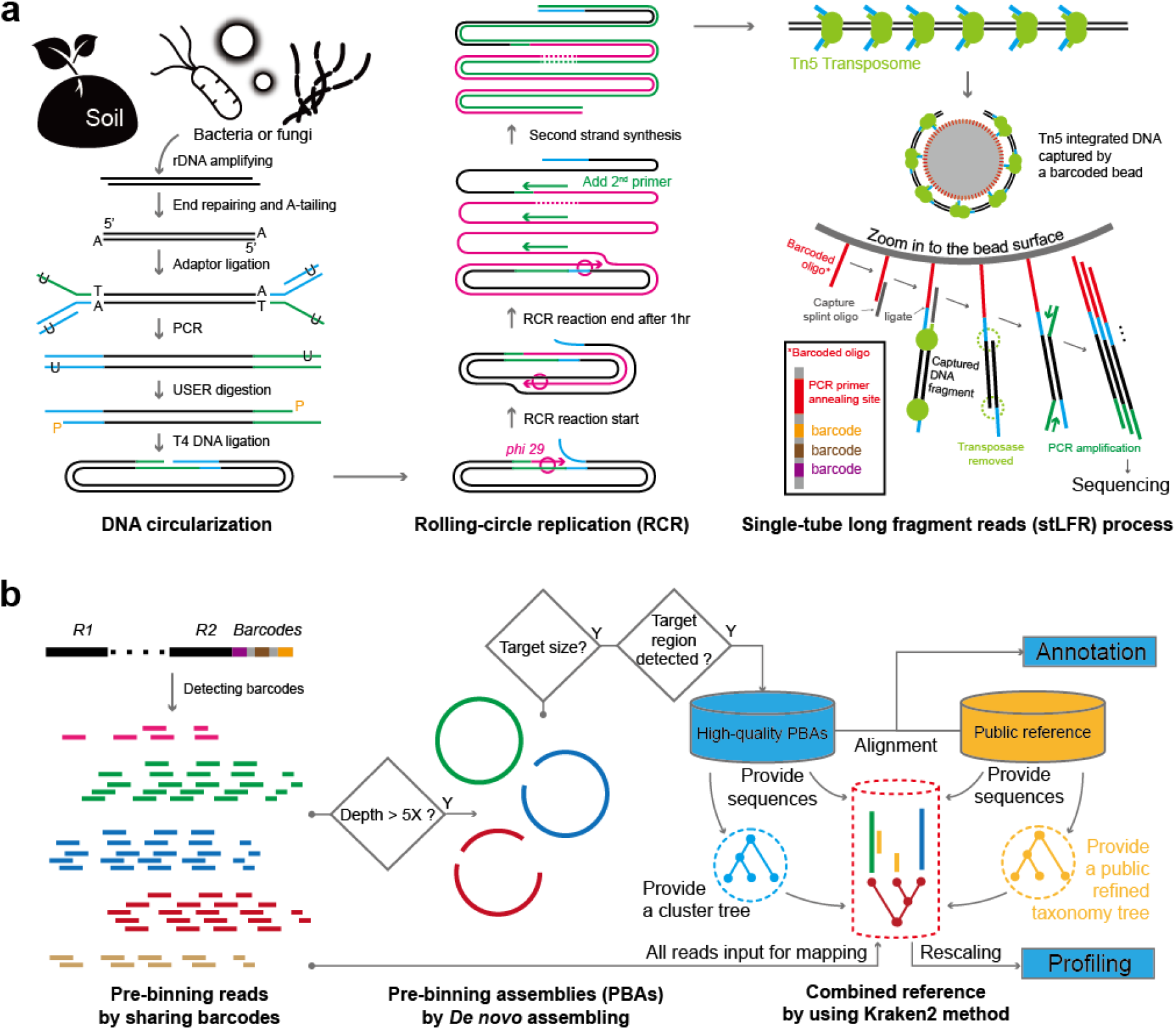
Sequencing framework (a). *In vitro* processes of rolling-circle replication and stLFR. DNA from bacteria and fungi was extracted and amplified using designed rDNA primers. Amplicons were turned into long concatemers with circularization and rolling-circle replication (RCR). The products of RCR were labeled with unique barcodes by integrating transposons and hybridizing onto 30 million different clonal barcoded beads in a single tube. After PCR, these sheared short sub-fragments were subjected to DNA sequencing. **(b)**. *In silico* processes of assembly, classification, and quantification. Reads with attached barcodes were decoded during the quality-control process. Pre-binning was done by grouping reads sharing the same barcode. Subsequently, *de novo* assembly was performed for each bin independently and in parallel. The pre-binning assemblies (PBAs) were then selected by target size range and were detected rDNA subunits or ITS regions by Barnnap and ITSx softweare were used for. The open reference databases SILVA and UNITE were used for taxonomic classification. Kraken2 was used to generate a database for profiling by mapping all barcode-detected reads.

The ZymoBIOMICS™ mock community which contains 8 bacterial species (See Methods) was used to test the ability of stcLFR to properly assemble the rDNA amplicons. A region covering 4.5 kb from the SSU (515Fng forward primer) to the LSU (TW13 reverse primer) was amplified for sequencing. From 186,580,349 clean reads generated from 3 replicates, we decoded 26,512,829 valid barcoded read bins of which 148,814 having a coverage > 5X were used for *de novo* assembly. With a relatively small number of reads and simple genomic content, the assembly proceeded faster and was more complete compared to general metagenomic assembly. We finally retrieved 146,509 assembled PBAs with a centroid length distribution close to 800 bp (Supplementary figure 1a). Most of them covered either the SSU (34.15%) or the LSU (38.58%). Only 25.23 % of PBAs covered both the SSU and the LSU (Supplementary figure 1b). The assembly quality was high, with 9.72% of the aligned PBAs exhibiting 100% identity to the reference genomes, and more than 60% of the PBAs achieved 99.5% or higher identity with the reference genomes. By contrast, only 1.78% of the PBAs could not align to the reference genomes (Figure 2a).

**Figure 2.**
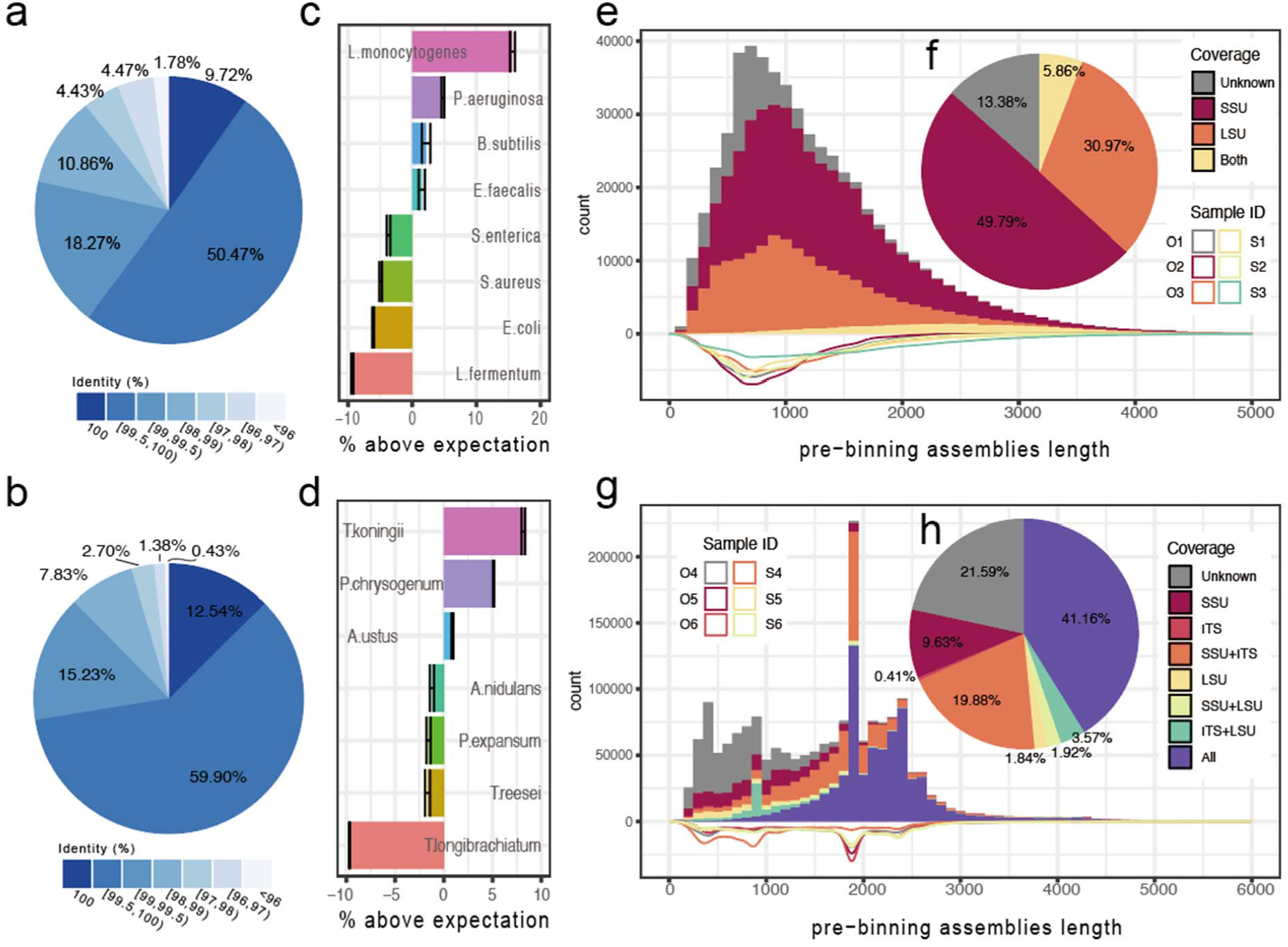
Long fragments restore performance. (a) Identity of alignments of mock bacterial rDNA PBAs to reference sequences. (b) Identity of alignments of mock fungal rDNA PBAs to reference sequences. (c) Difference between the observed and the calculated abundances of bacterial species in the mock community. (d) Difference between the observed and the calculated abundances fungal species in the mock community. (e) rDNA sequences assembled from soil bacterial DNA. Subunits (16S/SSU and 23S/LSU) were detected by alignment to the SILVA and UNITE database or predicting by barrnap. The length distribution of each sample is plotted on the negative part of the y-axis. Insert, pie plot of percentages of subunits detected. (f) rDNA sequences assembled from soil fungal DNA. Percentage of covered 18S/SSU, 28S/LSU and ITS is indicated by different colors. Insert, pie plot of percentages of subunits and ITS region detected.

For tests on fungi, a mock community comprising 7 common fungal species was used to assess the classification resolution at the species level (See Methods). Primers spanning a region ∼ 2.5 kb covering ITS1 and ITS2, and parts of the flanking SSU and LSU rRNA genes were used. Following the DNA library preparation and sequencing protocols used for bacterial species, we generated 190,208,124 high-quality reads, decoded 21,215,624 barcoded bins, with 270,794 having a coverage > 5X. In total, 263,416 PBAs were generated with a size centered on 2.3 kb, which is close to the value expected from the design (Supplementary figure 1c). Exceeding the results obtained for the bacterial samples, 12.54% of the fungal PBAs exhibited a 100% alignment to the reference genomes and more than 72.4% of the PBAs achieved 99.5% or higher identity to the reference genomes. Only 0.43% of the PBAs were unable to be aligned to the reference genomes (Figure 2b). 76.8% of the PBAs covered the entire region from the SSU to the LSU, and the ITS region was included in 83.7% of the PBAs (Supplementary figure 1d).

For assessing the accuracy of quantification, we mapped all decoded reads to the assembled complete rRNA gene sequences. Mapped reads sharing the same barcode represent a single DNA molecule and as such were counted once. We determined the difference between the observed and the calculated relative abundance of each species to estimate the error. For analyses of the mock bacterial community, we calculated the error to range from -9.3% to 15.6%, with an error standard deviation of 7.9% (Figure 2c). For analyses of the fungal mock community, we estimated the error to range from -9.7% to 8.2%, with an error standard deviation of 5.6% (Figure 2d). We estimated that the variation in the relative abundance of each species, as defined by the variance of error, was as low as 0.32% for bacterial species and 0.15% for fungi (Figure 2c). Though we only obtained a low percentage of amplicons covering the entire bacterial rRNA gene region as designed, we successfully recovered long fragments of bacterial 16S and 23S rRNA genes, and fungal fragments covering the entire ITS region and parts of the flanking 18S and 28S rRNA genes.

From both bacterial and fungal mock samples, the identity between PBAs and the corresponding reference sequences exceeded in most cases 99%, a percentage higher than 97% which often is used for amplicon-based species identification. In addition, the observed low variation in the relative abundance of each species also demonstrated the consistency of the protocol, an important requirement for comparative analyses using high-resolution profiles.

We had limited success in retrieving the entire ∼4 kb length of bacterial rDNA regions and observed a gap in the coverage of the bacterial rDNA genes, but still successfully recovered long fragments of bacterial 16S and 23S rRNA genes. The gap harbors transfer RNA (tRNA) genes (Supplementary figure 2), which may be a target for cleavage by the Tn5 transposase, the enzyme used for fragmentation during sequencing library preparation^16-19^. Although cleavage by the Tn5 transposase exhibits limited sequence bias, target preferences might still exist and cause the failure of effectively retrieving fragments covering the entire rDNA gene region^22-24^. In eukaryotes, tRNA genes are generally located outside the SSU and LSU regions, not in-between, which may at least in part explain why the eukaryotic amplicons survived, but the bacterial amplicons were split.

From the reference alignment benchmarking, we observed that when a PBA exceeded its designed size, part of the sequences could be aligned to reference sequences belonging to different species. We consider such PBAs as chimeric PBAs, which were subsequently filtered out (Supplementary figure 3b, red lines). Since we successfully enriched for the designed size of fungal rDNA sequences, this trend was prominent when the length of PBAs exceeded 2.3 kb for fungal samples. In addition, for bacterial PBAs with length < 500 bp we also observed a higher probability for achieving the exact same identity and bit score by multiple reference sequences representing different species, leading to an ambiguous annotation (Supplementary figure 3a, solid blue line). This trend became more severe in fungal PBAs with size < 2000 bp (Supplementary figure 3b, the solid blue line). These findings might point to one limitation of using short sequences to distinguish taxonomies at the species level, especially for fungi. However, coverage of multiple regions greatly eliminated the ambiguous annotation (Supplementary figure 4c).

To further explore the abundance and reproducibility of taxonomies in real environmental samples, we sequenced 3 replicates of two natural soil samples to identify bacterial rRNA genes and fungal ITS regions using the same primer sets and protocols used for the mock samples. We obtained 632,574,076 bacterial reads and 1,006,826,680 fungal reads with valid barcodes. For bacterial amplicons, we successfully generated 544,433 PBAs from 724,038 candidate bins (Figure 2e). As we observed using mock communities, only a small fraction (5.86%) of the bacterial PBAs covered both the SSU and the LSU regions. Most bacterial PBAs only covered the SSU (49.79%) or the LSU (30.97), whereas 13.38% did not match any currently known rDNA region (Figure 2f). For fungal amplicons, 1,807,779 PBAs from 1,845,429 high-coverage bins were generated with a size distribution that peaked around 2.3kb and 1.8kb (Figure 2g). A PBA size of 2.3kb was consistent with the primer design showing full coverage of the flanking region of the SSU region, the ITSs, and the flanking region of the LSU region. By contrast, half of the PBAs with sizes that peaked around 1.8kb lacked the LSU region. We detected ITS regions from 65.05% of the PBAs (Figure 2h). This percentage increased when the size exceeded 1 kb.

We also tried to profile the bacterial and fungal community at different rank levels. As soil samples contain very complex microbial communities, we chose Kraken2 to handle the profiling task. To build a Kraken2 database we first used PBAs to generate cluster trees for bacteria and eukaryotes (mostly fungi), respectively. Since we retrieved individual rDNA subunit assemblies for bacteria, we selected PBAs harboring SSU sequences for clustering. For eukaryotes, and especially fungi, the ITS region provided the highest level of discrimination, and accordingly, we selected PBAs covering ITS1 and ITS2 for clustering. The operational taxonomic unit (OTU) clusters were constructed using a set of identity thresholds enabling classification of taxa at multiple taxonomy levels, from domain to species. These thresholds were determined by pairwise global alignments of ribosomal gene subunit sequences including SSU and LSU from the SILVA database and ITS from the UNITE database (Supplementary figure 4), setting 97% identity as the threshold for genus association and 99% for species association for bacteria, and 95% identity and 97% identity as the thresholds for genus and species association, respectively, for fungi. Additional thresholds were the same for bacteria and fungi (See Methods). The trees generated using PBAs were then merged with taxonomy trees based on the SILVA and the UNITE databases (Figure 3a). In this process, clades were merged with taxonomies when the representing PBAs achieved sequence identities greater than the rank’s threshold. The combined tree contained three categories of branches. One type was represented by the public SILVA and UNITE sequences (PUBs, green in Figure 3, defined as exclusively PUBs), one represented by sequences only present in PBAs (blue in Figure 3, exclusively PBAs), and one represented by both the PBA and the PUB sequences (red in Figure 3, the shared PBAs and PUBs). The merged taxonomy tree and combined sequences were then used to build the database.

**Figure 3.**
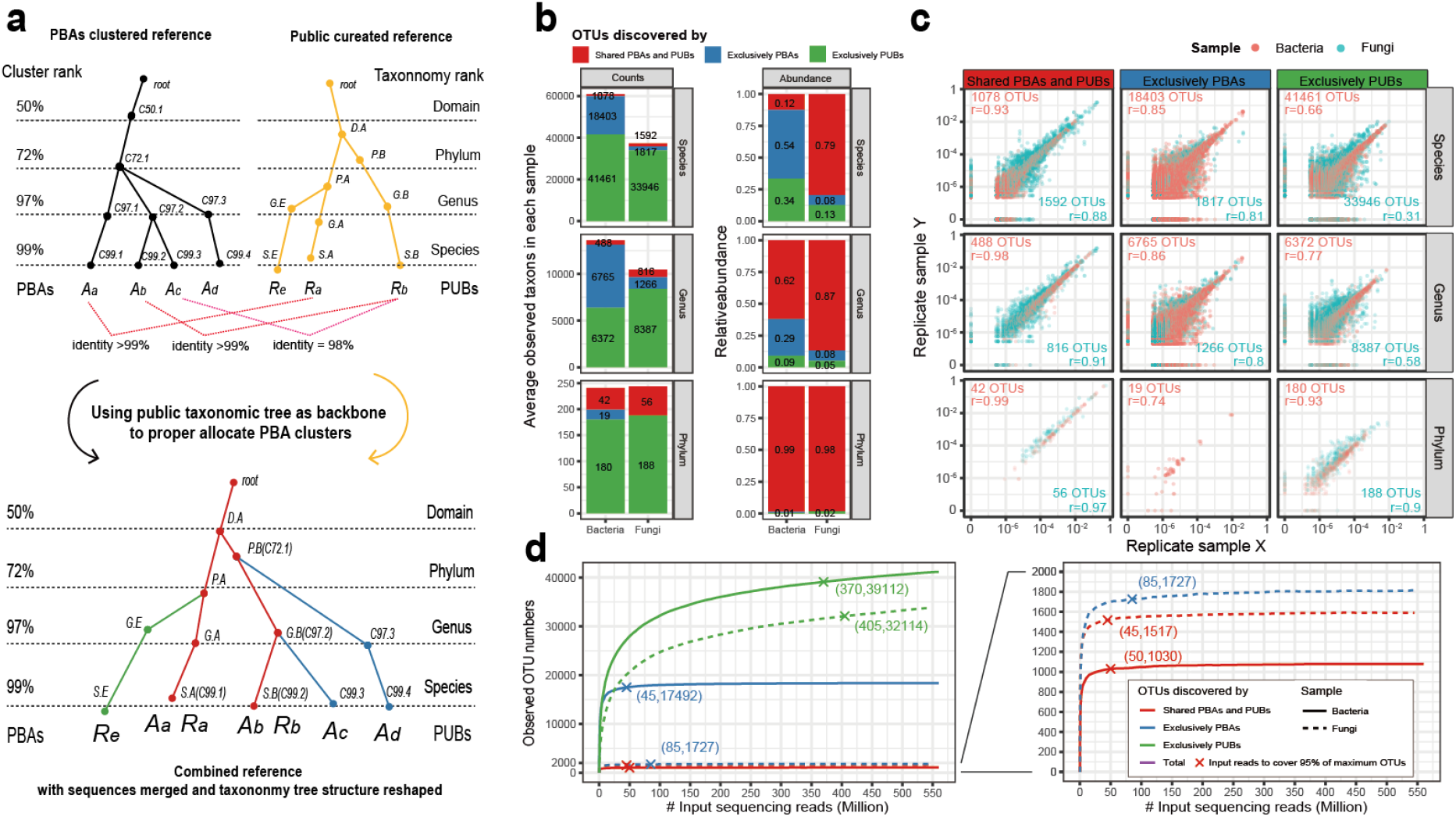
Identification and performance of a combined Kraken2 profiling strategy. (a) The pre binning assembly (PBA) cluster tree was generated by a series of clustering of PBAs at default identity cut-off for each level of taxonomy. The identity cut-offs were estimated by pairwise alignments of annotated SSU, LSU and ITS sequences from public databases. The public taxonomy tree was mainly based on the SILVA SSU taxonomy tree and supplemented with new taxonomies from the SILVA LSU and UNITE databases. A combination of PBAs and public reference sequences (PUBs) was established based on the identities between the PBA and PUB sequences. The combined taxonomy tree contains three branch categories of sequences: 1) Sequences shared by PBAs and the PUBs (red); 2) sequences unique to PBAs with no taxonomy association (blue); 3) Reference sequences from public taxonomy tree not shared by PBA (green). (b) The performance of the three branches of the taxonomic trees. Left: The number of OTUs shown at the species, genus and phylum levels, visualized as bar plots. Right, relative abundances. The colors indicate OTUs belonging to PUBs (green) or PBAs with (red) or without (blue) annotation. (c) Correlation of relative abundance between duplicates. At each level of taxonomy, the relative abundance observed in each duplicate is colored light red for bacteria and light green for fungi. The Spearman correlation values for bacteria and fungi are indicated (d) Rarefaction curves of observed bacterial and fungal OTUs at the species level using the different databases. One bacterial sample (solid line) and one fungal sample (dash line) with 3 replicates and sequenced to more than 500 million read pairs were analyzed. On each curve, a cross marks the number of read needed to cover 95% of the total counts.

By this approach, we mapped all barcoded reads to the combined database retrieving 60,942 bacterial OTUs at the species level of which 1,078 OTUs were supported by both the PBA sequences and PUB sequences, 18,403 could not be annotated, representing putative novel species, and the remaining 41,461 were annotated by publicly available sequences. At the genus level, 6,765 OTUs represented PBAs with no annotation. Using 72% identity as the threshold for association with a phylum, we identified 241 OTUs of which 19 were novel. For the eukaryotic communities, we retrieved 37,355 fungal OTUs at the species level of which 1592 OTUs were supported by both the PBA sequences and PUB sequences, 1,817 could not be annotated, representing putative novel species, and the remaining 33,946 OTUs were annotated by PUB sequences. At the genus level, 1,266 OTUs represented PBAs with no annotation. Using 72% identity as the threshold for association with a phylum, we identified 244 OTUs, all of which were annotated. (data of sample S are shown, Figure 3b, left panel).

We found that species level OTUs supported by both PBA and PUB sequences represented only 1.8% of all OTUs; the relative abundance of these OTUs combined corresponded to about 12% of the relative abundances of all OTUs. At the genus level, the relative abundance of OTUs supported by both the PBA and PUB sequences was even higher (62%), and at the phylum level the relative abundance of the OTUs supported by both PBA and PUB sequences reached close to 100%. This trend was more obvious for fungal OTUs, where 4.2% OTUs supported by both the PBAs and PUBs contributed 79% of relative abundance at the species level (figure 3b, right panel). This indicated that OTUs supported by PBAs (whether annotated or novel) were abundant and dominated the microbial community. Compared with the fungal community, more dominant taxonomies may represent novel species.

Further analysis showed that the three categories of branches in the combined tree exhibited distinct differences in terms of reproducibility between replicates. OTUs supported by both PBA and PUB sequences exhibited high correlation between replicates (r=0.93 for bacterial communities and r=0.88 for eukaryotic communities; Spearman’s correlation, figure 3c). Higher correlations were observed at the genus and the phylum levels. By contrast, the correlation between replicates of OTUs supported exclusively by PBA was lower and the correlation decreased significantly when the abundance became lower than 10^−4^, both for bacteria and eukaryotes. Some of the unannotated bacterial phyla taxa still varied, indicating that they might not represent an independent phylum. For the taxa supported by public references, though well organized by taxonomic information, the species taxa performed worse than those supported by unannotated PBAs. However, the performance was improved dramatically at the phylum levels. These dominant annotated OTUs supported both by PBAs and PUBs and highly abundant (>10^−4^) novel OTUs exhibited high reproducibility for quantification comparison tasks at the species and the genus level.

The rarefaction curves also showed a better performance in terms of number of reads needed for sufficient coverage of OTUs from the shared PBAs and PUBs sequences. To retrieve the majority of taxa (here defined as 95% of maximum OTUs), the publicly available database required the highest number of read-pairs, 410 million read-pairs for bacteria and 425 million read-pairs for fungi. To assess exclusively PBA OTUs, this requirement was reduced to 110 million and 150 million for bacteria and eukaryotes, respectively. The shared PBAs and PUBs representing OTUs required the lowest number of reads, only 40 million bacterial read-pairs and 55 million eukaryotic read-pairs, respectively.

An alternative approach for long-read amplicon sequencing was recently published by Karst and co-workers^20^. In this work, re-designed UMIs and single-molecular sequencing were used for obtaining long-read high-accuracy amplicon sequences using Nanopore or PacBio sequencing. While this is an elegant approach, our approach relying on second-generation sequencing enables a more cost-effective very high sequencing throughput coupled with robust and reproducible quantification. In summary, our approach provided a 99% identity of species level, high-throughput strategy to expand current rRNA gene databases by including long marker sequences and potential novel taxonomies, as well as a comparable accurate quantification profiling strategy. This approach provides a cost-effective method for obtaining extensive and accurate information on environmental complex microbial communities.

## Methods

### Sample collection and DNA extraction

The mock bacterial DNA standard, ZymoBIOMICS Microbial Community Standard (D6305), was purchased from ZymoBIOMICS™. Fungi strains were purchased from China General Microbiological Culture Collection Center. The fungal mock DNA sample consisted of equally mixed genomic DNA of 7 fungi, including *Aspergillus ustus* (No. BNCC144426), *Trichoderma koningii* (No. BNCC144774), *Penicillium expansum* (No. BNCC146144), *Aspergillus nidulans* (No. BNCC336164), *Penicillium chrysogenum* (No. BNCC336234), *Trichoderma reesei* (No. BNCC341839) and *Trichoderma longibrachiatum* (No. BNCC336352). The sequence of the rDNA region for each strain was rechecked by Sanger sequencing. Soil samples were collected from Nanbanhe tropical rainforest plot (21.612 N, 101.574 E) in Yunnan, China in 2017. Soil samples were immediately stored at -80°C after collection, and DNA extraction was performed within two months. DNA extraction of all samples was performed using the PowerSoil®DNA Isolation Kit (Mobio) according to manufacturer’s instructions. DNA concentration was measured by Qubit flex fluorometer (Invitrogen).

### rDNA long fragments amplification

We initially amplified the region of bacterial 16S-23S rRNA gene and fungi full length ITS region with Kapa Hifi DNA polymerase (Roche) and EX Taq DNA polymerase (Takara Bio). No correct PCR products (examined by Sanger sequencing) were amplified by Kapa, probably due to its high fidelity and the fact that the conserved regions where primers locate can result in mismatches of several nucleotides. By contrast, EX Taq DNA polymerase (Takara Bio), which also has proofreading activity, successfully amplified the rDNA targets, it was therefore used for full length rDNA amplification. For PCR amplification we used 5’ phosphorylated primers. The primer sequences are listed below. For bacteria, the forward primer was 27F (AGAGTTTGATCATGGCTCAG) and the reverse primer was our in-house designed 23S-2850R (CTTAGATGCCTTCAGCRVTTATC). For fungi, highly universal eukaryotic primers for high eukaryotic and fungal taxonomic coverage were selected based on a previous study (Tedersoo et al., 2018). The forward primer was SSU515Fngs (GCCAGCAACCGCGGTAA) and the reverse primer was TW13 (GGTCCGTGTTTCAAGACG). The conditions for PCR were as follows: 1 µL of diluted template DNA, 1 µL of forward primer (10 M), 1 µL of reverse primer (10 M), 15.5 µL of nuclease-free water, 5 µL of 10X Ex Taq Buffer, 1 µL of dNTPs Mixture (2.5 mM each), and 0.5 µL of EX Taq Polymerase. We amplified samples using the following cycling conditions: 95 °C for 5 min; 30 cycles of 95°C for 30s, 55°C for 30s, and 72°C for 3 min; and then a final extension at 72 °C for 10 min. The amplified long amplicons were purified using QIAquick PCR Purification Kit (Qiagen).

### Circularization and Rolling Circle Replication

To enable a highly efficient circularization of long molecules, a double stranded circularization method was applied. 100 ng amplicon were first subjected to end-repair and A-tailing according to the protocol described previously^21^. Then the product was incubated with 8 pmol of adapter and 3000 units T4 DNA ligase (MGI, 1000004279) in 80 μL of 1X PNK buffer (NEB, B0201S) with extra 1 mM ATP and 7.5% PEG-8000 at room temperature for 1 hour, followed by a 0.5X SPRI beads purification (Beckman, A63882) and eluted with 20 μL of TE buffer (Thermo Fisher, AM9849). Next polymerase extension with 1 pmol of primer containing uracil was carried out, by adding prior heat-activated 200 units Pfu Turbo Cx (Agilent Technologies, Inc., 600414) in 50 μL of 1X PfuCx buffer at 72°C for 20 minutes, followed by a 1.5X SPRI beads purification and elution with 30 μL of TE buffer. Sticky ends were created by injecting 20 units of USER enzyme (NEB, M5505S) and incubating with 50 μL of 1X TA buffer (Teknova, T0380) at 37°C for 1 hour. T4 DNA ligase-mediated circularization was performed in 150 μL of 1X TA buffer with extra 1 mM ATP. All linearized DNA were removed with 0.4 units Plasmid-Safe™ DNase (Lucigen, E3110K) followed by a 1X SPRI beads purification.

The double stranded circle from the last step was designed with a 3nt gap on one strand acting as the initial extending site for rolling circle replication. The RCR reaction was carried out by incubation with 5 units Phi29 (MGI, 1000007887) in 21 μL of 1X Phi buffer at 30°C for 1 hour. In this step the original long amplicons were transformed into long concatemers with multiple copies of each long molecule. Next, unlike other studies^22, 23^, we applied 60 units warm-started Bst 2.0 polymerase (NEB, M0538M) and primer extension with specific sequences (CGCTGATAAGGTCGCCATGCCTCTCAGTAC) to generate the second strand of the RCR product ready for standard stLFR library preparation.

### stLFR barcoded DNA library construction

Extended amplicons produced after RCR were labeled by unique barcodes with MGIEasy stLFR Library Prep Kit (MGI, 1000005622). Briefly, indexed transposons are inserted into 1ng of double strand rolling circle replication products from different samples, followed by hybridization of the transposon integrated DNA onto clonally barcoded beads. After capture, the sub-fragments of each transposon inserted DNA molecule are ligated to the barcode oligo. A few additional library processing steps are performed followed by PCR. At this point the co-barcoded sub-fragments are ready for sequencing. This method and a detailed protocol were previously described by Wang et. al.^24^ and Cheng et. al.^25^, respectively.

### Sequencing and decoding long fragments

DNA libraries were sequenced using the pair-end 100 bp mode on the BGISEQ-500 platform^26^. During sequencing, the barcode part was sequenced first and attached to the tail of read2. The barcode detection and sequence read quality control were managed by the fastp software^27^.This tool was initially designed for traditional NGS data QC, with a parallel function to speed up the process. We added barcode detection module to it. In the detection process, each 10-base barcode sequence (3 10-base sequences make up a full barcode) is scanned against our available barcode list, both by forward and reverse strand, within 1 base mismatch tolerance. In addition, a module to perform the BGISEQ platform specific quality filtering and trimming was also implemented as described previously^26^. Reads were sorted by barcode and stored in a fastq format. More than 85% of reads were associated with a valid barcode. Less than 1% of detected barcodes were recovered after barcode error correction (data not shown).

### Single-molecular pre-binning and *de novo* assembly strategy

An estimation of kmer coverage was initially performed to ensure that each bead had enough coverage for an independent de novo assembly. The kmer coverage was determined by the formula:

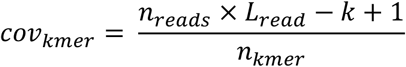

Where *n*_reads_ means reads number belonging to a bead; *L*_read_ means the length of each read, which is 100bp here; *k* means the length of kmer, which is 31; *n*_kmer_ means the unique number of kmers from *n*_reads_, here calculated by mash version 2.1.1^28^.

According to the coverage distribution, we only allowed beads with *cov*_kmer_≥5 for the assembly process. Each bead’s assembly was performed by megahit version 1.1.2 ^29^, with parameters *--k-min 21 --k-step 20 --prune-level 0 --min-count 1*.

Since amplicons were head to tail adjoined by the RCR adaptor, it is possibile to assemble a circular sequence with the RCR adaptor. In this case, a script to clip the RCR adaptor from contigs and linearize the sequence was used.

### Taxonomic annotation for PBAs

For taxonomic assignment, PBAs were initially aligned to SILVA version 138 SSU and LSU refseq^30^ by blastn^31^, separately. For fungal PBAs, UNITE^32, 33^ (released at Apr 2, 2020) was used as reference of ITS region. Alignment results were then summarized to provide the most possible taxonomy for each PBA according to the summary of alignments of the SSU, ITS and LSU regions, measured by sequence identity, bit score, and length coverage. PBAs with ambiguous annotation pointing to multiple different taxonomies with the same score were eliminated. Chimeric PBAs with several fragments uniquely assigned to different taxonomies were also discarded.

### Default taxonomic rank threshold determination

The pairwise global alignments were performed for the SSU, LSU and ITS region sequences, separately. The SSU sequences were collected from SILVA version 138.1 SSU refseq^30^. The LSU sequences were collected from SILVA version 138.1 LSU refseq^30^. The ITS sequences were collected from UNITE^32, 33^ (released on Apr 2, 2020). For each region, the alignments were initiated from its top rank (species or subspecies). For each taxonomy, if multiple associated sequences existed, a pairwise global alignment was performed by vsearch v2.14.1^34^ to calculate sequences identities between each two sequences, with the following command:

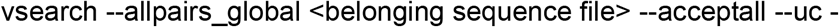

To minimize computing, no more than 100 sequences from a given taxa were used. After completion of assignment to all ranks, the sequence identity distribution of the median value of each taxon was used for visualization. For each rank, the peak position was selected manually as the estimated threshold for this rank.

### Cluster tree generation

For each rank, taxonomy assignment of a given taxon was determined and organized for Kraken version 2.0.8-beta^35^ database generation.

Bacterial PBAs ranging from 500 bp to 1500 bp and fungal PBAs ranged from 1700 bp to 2500bp were picked

PBAs used to process clusters satisfied the following criteria: 1) target rDNA region(s) detected; 2) size of length in defined range; 3) no ambiguous annotation; 4) no chimera detected. For bacterial PBAs, barrnap version 0.9 (https://github.com/tseemann/barrnap) was used for rDNA regions detection, and the picked length ranged from 500 bp to 1500bp. For fungal PBAs, barnnap version 0.9 and ITSx version 1.0.11^36^ were both used for rDNA regions detection, and picked length ranged from 1700 bp to 2500 bp. Any PBA with partial segments aligned with conflicting annotations was discarded. Retained PBAs were then clustered by vsearch with a series of identities at 0.4, 0.5, 0.66, 0.72 (for phylum), 0.77, 0.83, 0.89, 0.92, 0.95, 0.97(for genus), 0.98, 0.99 (for species), 0.995, 0.999 and 1, according to the taxonomies similarity centroids calculated from all annotated sequences from the SILVA and the UNITE databases (See Supplemental table 3). Each cluster with a certain identity was regard as a clade of a taxonomy tree if the member PBAs could not provide a unified rank annotation. Note that the cutoff of rank above species varied greatly among different groups. The cutoff values were only assigned for clades without any useful information, otherwise they were determined by the clade members’ classifications. Finally, at species rank, singleton OTUs without annotation were discarded in order to reduce the false discovery rate.

### Relative abundance computation of each rank

The rank abundance was calculated by Kraken2, which uses kmer alignments to determine the position of each read. We then rescaled it to barcode unit whereby reads sharing the same barcode represent a single DNA molecule. The generated results are compatible with Kraken2 so each rank’s profile can be classified directly.

### Phylogenetic tree building

For bacterial SSU rRNA gene sequences, barrnap version 0.9 (https://github.com/tseemann/barrnap) was used to secure quality of the PBAs sequences with no chimeras. Fungal ITS regions were predicted by ITSx version 1.0.11^36^ which also enabled determination of the boundary of the SSU and the LSU. The SSU and LSU sequence were then joined together for heterogeneity rate computation. Both bacterial SSU and fungal joint SSU+LSU sequences were multiple aligned by mafft v7.407^37^ and terminal gaps trimmed by trimAl v1.4.rev22^38^. RAxML version 8.2.1^39^ was then employed to perform 100 boostraps General Time Reversible model of nucleotide substitution under the Gamma model of rate heterogeneity, with accommodated searches incorporated (GTRCAT). The processes were executed on a 40-core node of the Danish National Supercomputer for Life Sciences (Computerome 2.0) with following parameters:

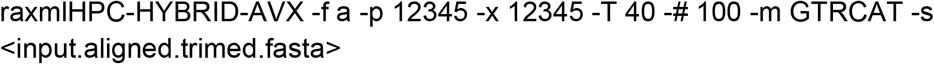

The results of the best-scoring ML tree with support values were imported and visualized by ggtree package^40^ in Rstudio v1.3.1073^41^ IDE for R 4.0.3^42^.

### Data availability

The data that support the findings of this study have been deposited into CNGB Sequence Archive (CNSA)^43^ of China National GeneBank DataBase (CNGBdb)^44^ with accession number CNP0001509. All data processing scripts are integrated in a toolbox pipeline available on github (https://github.com/Scelta/metaSeq).

## Acknowledgements

Ou Wang is a recipient of and this work was partially supported by National Natural Science Foundation of China (32001054). The samples were obtained from Dr. Yuehua Hu, who is funded by the West Light Foundation of the Chinese Academy of Sciences to Yue-Hua Hu.

## Contributions

F.F, O.W. and B.P. conceived the wet lab method. X.S., F.F. and O.W performed wet lab experiments and BGISEQ500 sequencing. C.F. and Z.S. conceived the bioinformatics method. C.F developed the software pipeline and performed data analysis as well as visualization. X.S. and X.Z. interpreted the microbial characteristics of data. X.S. performed the phylogenetic analysis. H.Z., Z.P. and X.L. assisted to perform the data. C.F., X.S., X.Z., O.W., B.P., Z.S. and K.K. wrote the manuscript. All authors participated in discussions and contributed to the revision of the manuscript. All authors read and approved the final manuscript.

## Supplementary Material

**Supplementary figure 1.**
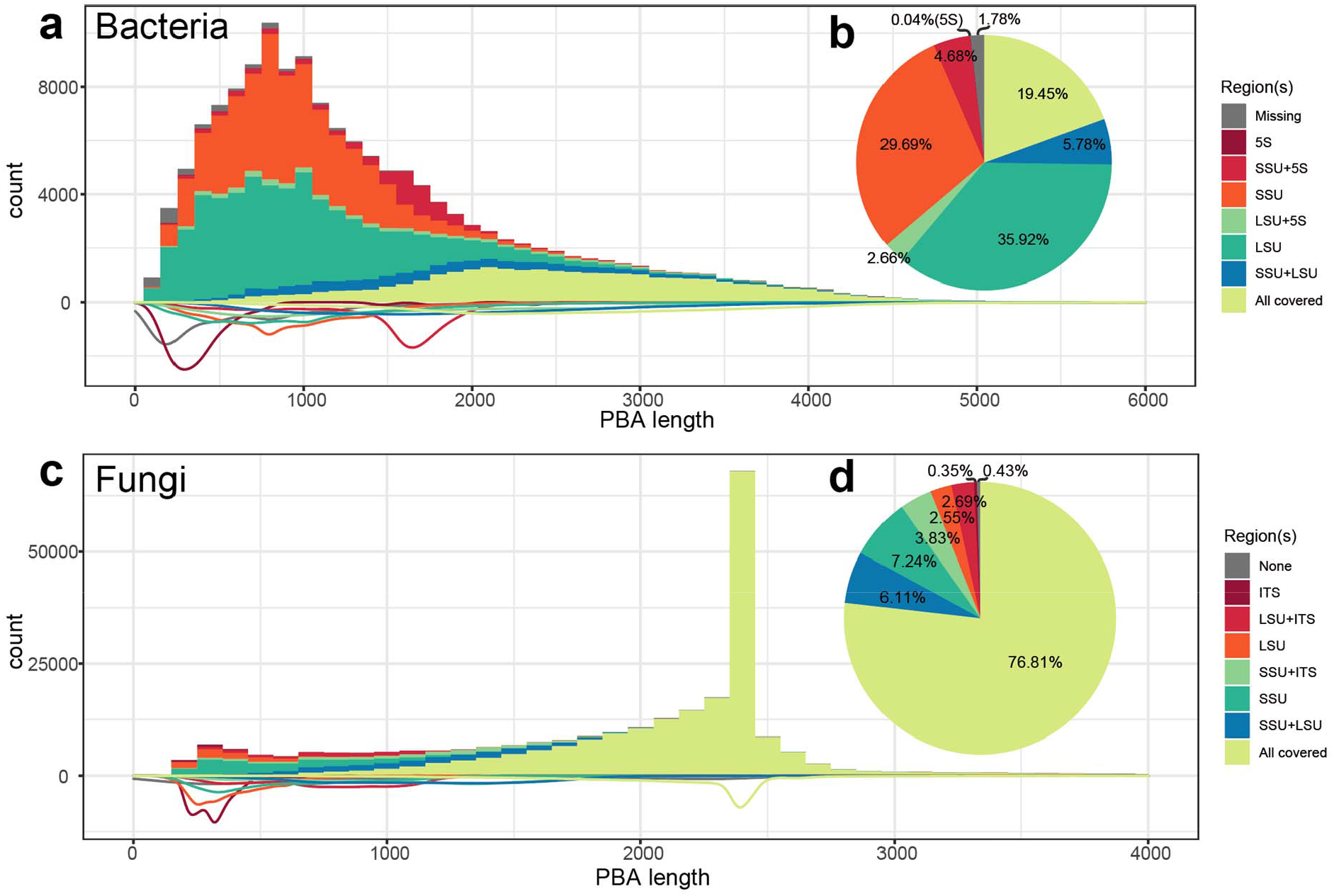
Length and rDNA subunits coverage of bacterial and fungal PBAs. (a) Length distribution of bacterial mock PBAs. Percentages of SSU, LSU and 5S coverage are shown in colors. Density lines of subunit coverage are shown on the negative side of the y-axis. (b) Pie plot of percentage of subunits detected. (c) Length distribution of fungal mock PBAs. Percentages of SSU, LSU and ITS coverage are shown in colors. Density lines of subunit coverage conditions shown on the negative side of the y-axis. (d) Pie plot of percentages of subunits and ITS region detected.

**Supplementary figure 2.**
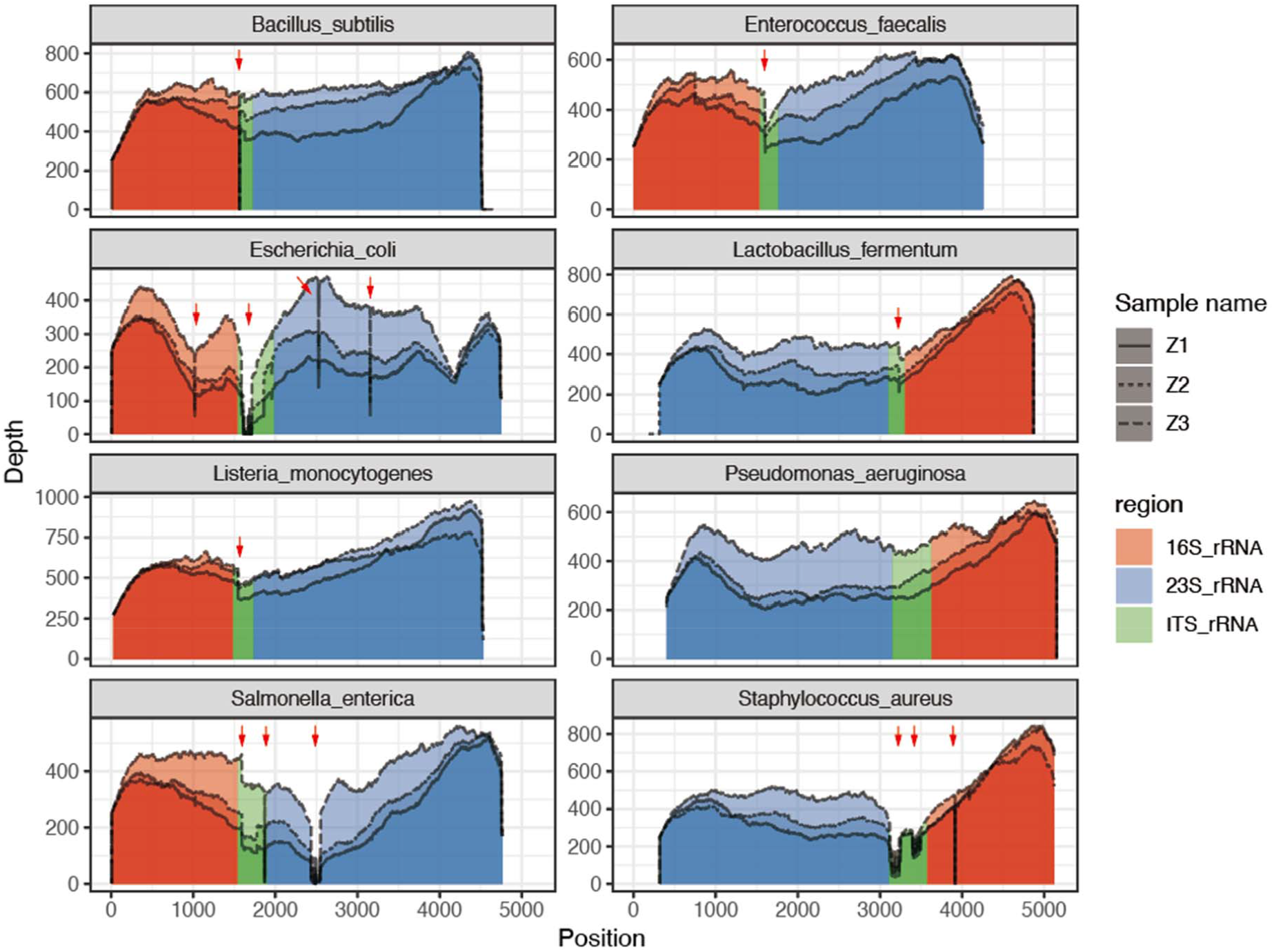
PBA coverage of 8 bacterial mock species’ rRNA gene sequences. All PBAs were mapped to their corresponding reference sequences. 16S, ITS and 23S regions were detected by barrnap and colored. Several positions where the coverage immediately dropped were observed in some species (red arrows).

**Supplementary figure 3.**
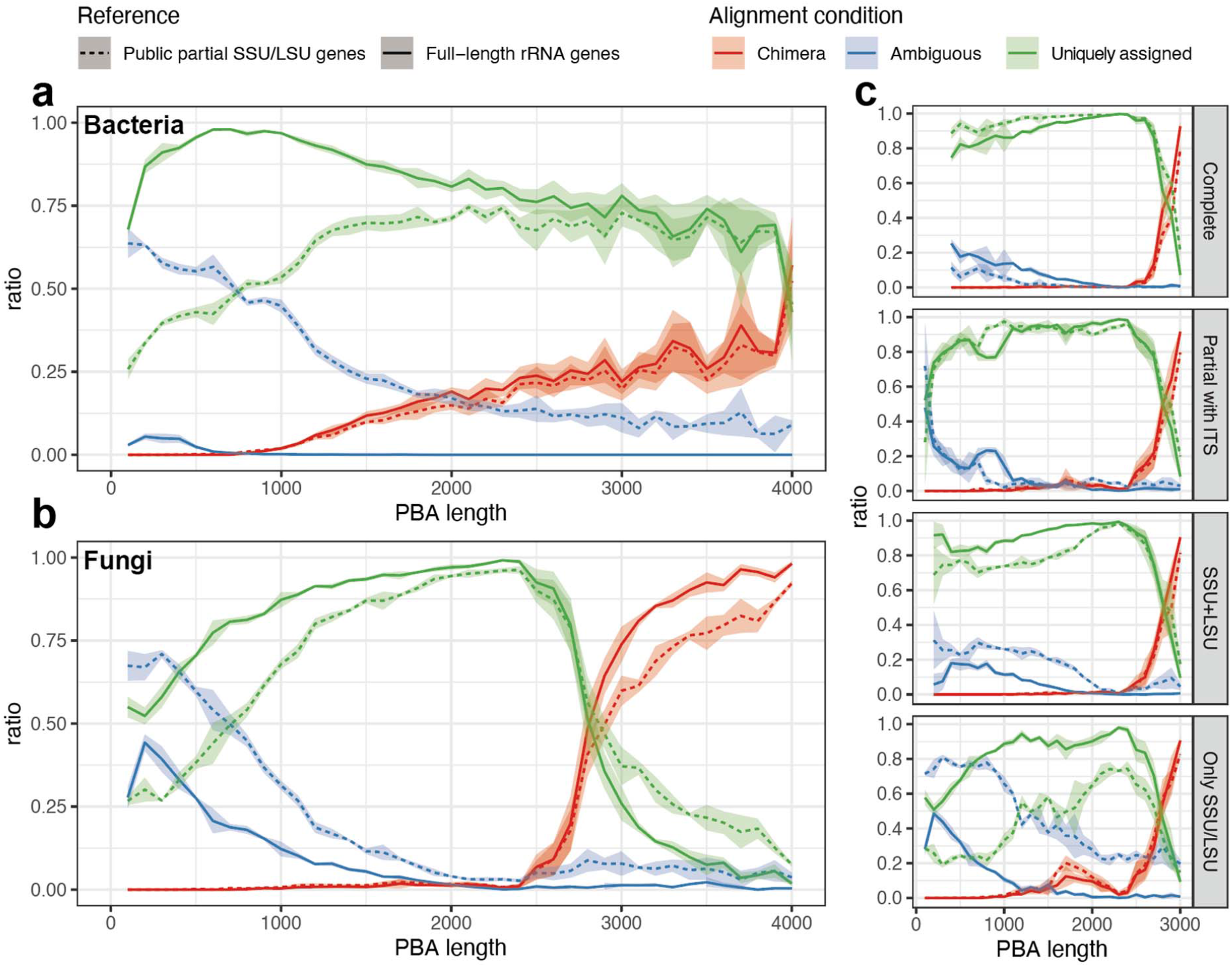
Taxonomic classification accuracy estimation. Fraction of ambiguous (blue) and chimera PBAs (red) were summarized in each 100 bp window of PBA length in bacterial (a) and fungi (b) samples, respectively. Mock specific complete reference sequences (solid line) and public references from PUB (dashed line) were used for comparison. (c) Fungal PBAs were divided according to covered regions for estimation of the ratio of ambiguous and chimeric sequences relative to the length of the PBAs..

**Supplementary figure 4.**
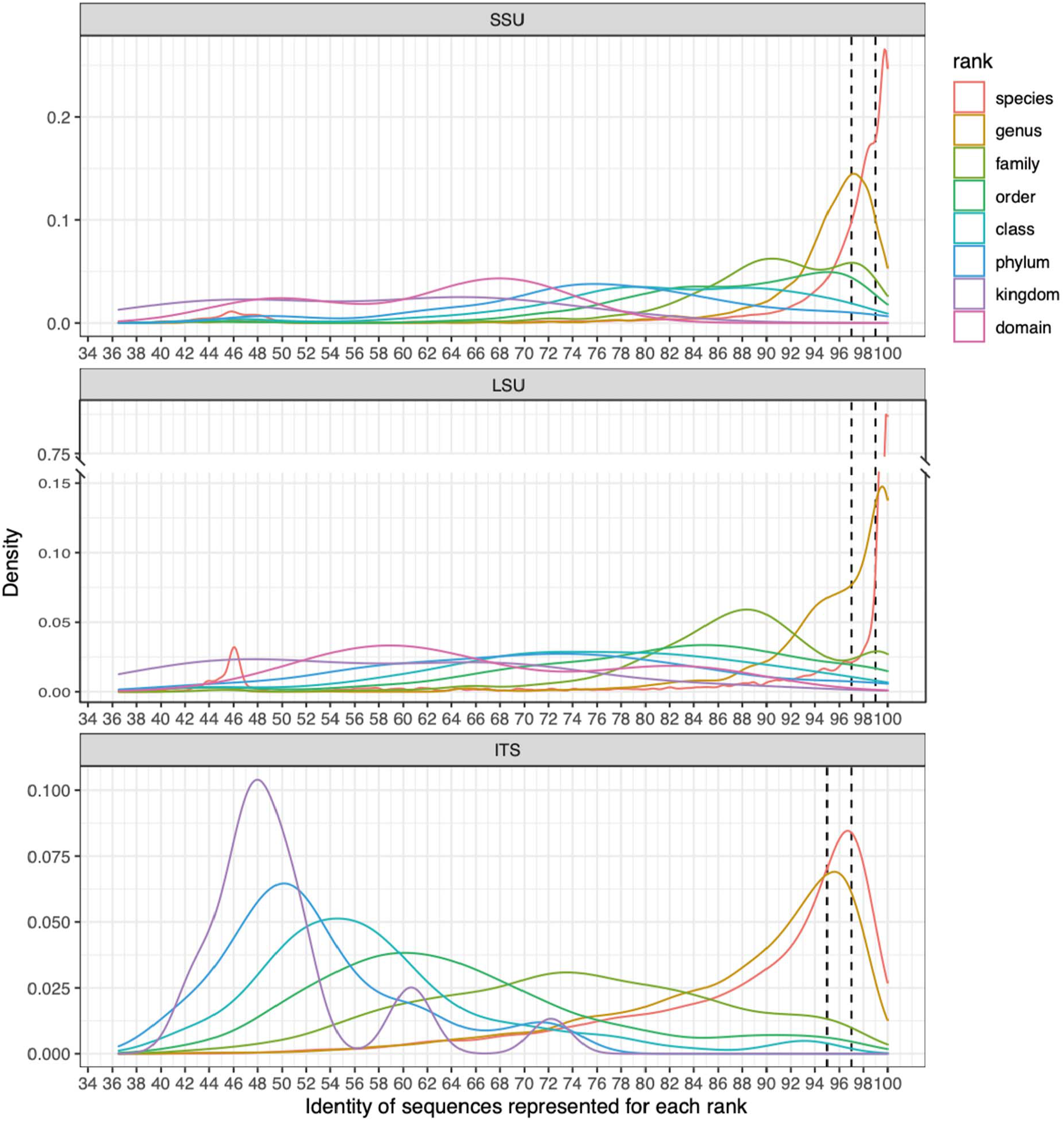
The sequence identity distribution of each rank based on rRNA gene sub-region sequences. The median value of sequence identities between each two members of a given taxonomy was used. At each rank, the curve peak position was selected manually as the estimated threshold to determine a default rank of an unknown clade.

## Reference

1. Vogel, T.M. et al. TerraGenome: a consortium for the sequencing of a soil metagenome. Nature Reviews Microbiology 7, 252–252 (2009).

2. Ehrlich, S.D. in Metagenomics of the Human Body. (ed. K.E. Nelson) 307–316 (Springer New York, New York, NY; 2011).

3. Gevers, D. et al. The Human Microbiome Project: a community resource for the healthy human microbiome. PLoS Biol 10, e1001377 (2012).

4. Gilbert, J.A., Jansson, J.K. & Knight, R. The Earth Microbiome project: successes and aspirations. BMC biology 12, 69 (2014).

5. Sunagawa, S. et al. Structure and function of the global ocean microbiome. Science 348, 1261359 (2015).

6. Bahram, M. et al. Structure and function of the global topsoil microbiome. Nature 560, 233–237 (2018).

7. Hamady, M., Lozupone, C. & Knight, R. Fast UniFrac: facilitating high-throughput phylogenetic analyses of microbial communities including analysis of pyrosequencing and PhyloChip data. The ISME Journal 4, 17–27 (2010).

8. Schoch, C.L. et al. Nuclear ribosomal internal transcribed spacer (ITS) region as a universal DNA barcode marker for <em>Fungi</em>. Proceedings of the National Academy of Sciences 109, 6241–6246 (2012).

9. Bishara, A. et al. High-quality genome sequences of uncultured microbes by assembly of read clouds. Nature biotechnology (2018).

10. Karst, S.M. et al. Retrieval of a million high-quality, full-length microbial 16S and 18S rRNA gene sequences without primer bias. Nature biotechnology 36, 190–195 (2018).

11. Deamer, D., Akeson, M. & Branton, D. Three decades of nanopore sequencing. Nature biotechnology 34, 518–524 (2016).

12. Wagner, J. et al. Evaluation of PacBio sequencing for full-length bacterial 16S rRNA gene classification. BMC Microbiology 16, 274 (2016).

13. Benítez-Páez, A., Portune, K.J. & Sanz, Y. Species-level resolution of 16S rRNA gene amplicons sequenced through the MinION™ portable nanopore sequencer. GigaScience 5 (2016).

14. Peters, B.A., Liu, J. & Drmanac, R. Co-barcoded sequence reads from long DNA fragments: a cost-effective solution for “perfect genome” sequencing. Frontiers in Genetics 5 (2015).

15. Wang, O. et al. Efficient and unique cobarcoding of second-generation sequencing reads from long DNA molecules enabling cost-effective and accurate sequencing, haplotyping, and de novo assembly. Genome research 29, 798–808 (2019).

16. Adey, W. Algal Turf Scrubber (ATS), Algae to Energy Project: Cleaning Rivers while Producing Biofuels and Agricultural and Health Products. Progress Report to the Lewis Foundation. Smithsonian Institution (2010).

17. Picelli, S. et al. Tn5 transposase and tagmentation procedures for massively scaled sequencing projects. Genome research 24, 2033–2040 (2014).

18. Hennig, B.P. et al. Large-scale low-cost NGS library preparation using a robust Tn5 purification and tagmentation protocol. G3: Genes, Genomes, Genetics 8, 79–89 (2018).

19. Wang, Y. et al. A practical random mutagenesis system for Ralstonia solanacearum strains causing bacterial wilt of Pogostemon cablin using Tn5 transposon. World Journal of Microbiology and Biotechnology 35, 7 (2019).

20. Karst, S.M. et al. High-accuracy long-read amplicon sequences using unique molecular identifiers with Nanopore or PacBio sequencing. Nature methods 18, 165–169 (2021).

21. Dong, Z. et al. Development of coupling controlled polymerizations by adapter-ligation in mate-pair sequencing for detection of various genomic variants in one single assay. DNA Research 26, 313–325 (2019).

22. Volden, R. et al. Improving nanopore read accuracy with the R2C2 method enables the sequencing of highly multiplexed full-length single-cell cDNA. Proceedings of the National Academy of Sciences 115, 9726–9731 (2018).

23. Adams, M. et al. One fly–one genome: chromosome-scale genome assembly of a single outbred Drosophila melanogaster. Nucleic acids research 48, e75–e75 (2020).

24. Wang, O. et al. Efficient and unique cobarcoding of second-generation sequencing reads from long DNA molecules enabling cost-effective and accurate sequencing, haplotyping, and de novo assembly. Genome research 29, 798–808 (2019).

25. Peters, B. et al. A simple bead-based method for generating cost-effective co-barcoded sequence reads. Protocol Exchange (2018).

26. Fang, C. et al. Assessment of the cPAS-based BGISEQ-500 platform for metagenomic sequencing. Gigascience 7, 1–8 (2018).

27. Chen, S., Zhou, Y., Chen, Y. & Gu, J. fastp: an ultra-fast all-in-one FASTQ preprocessor. Bioinformatics 34, i884–i890 (2018).

28. Ondov, B.D. et al. Mash: fast genome and metagenome distance estimation using MinHash. Genome biology 17, 132 (2016).

29. Li, D., Liu, C.M., Luo, R., Sadakane, K. & Lam, T.W. MEGAHIT: an ultra-fast single-node solution for large and complex metagenomics assembly via succinct de Bruijn graph. Bioinformatics 31, 1674–1676 (2015).

30. Quast, C. et al. The SILVA ribosomal RNA gene database project: improved data processing and web-based tools. Nucleic acids research 41, D590–D596 (2012).

31. Altschul, S.F., Gish, W., Miller, W., Myers, E.W. & Lipman, D.J. Basic local alignment search tool. J Mol Biol 215, 403–410 (1990).

32. Nilsson, R.H. et al. The UNITE database for molecular identification of fungi: handling dark taxa and parallel taxonomic classifications. Nucleic acids research 47, D259–D264 (2019).

33. Abarenkov, K.Z., Allan; Piirmann, Timo; Pöhönen, Raivo; Ivanov, Filipp; Nilsson, R. Henrik; Kõljalg UNITE general FASTA release for eukaryotes 2. Version 04.02.2020.. UNITE Community (2020).

34. Rognes, T., Flouri, T., Nichols, B., Quince, C. & Mahe, F. VSEARCH: a versatile open source tool for metagenomics. PeerJ 4, e2584 (2016).

35. Wood, D.E., Lu, J. & Langmead, B. Improved metagenomic analysis with Kraken 2. Genome biology 20, 257 (2019).

36. Bengtsson-Palme, J. et al. Improved software detection and extraction of ITS1 and ITS2 from ribosomal ITS sequences of fungi and other eukaryotes for analysis of environmental sequencing data. Methods in Ecology and Evolution, n/a-n/a (2013).

37. Nakamura, T., Yamada, K.D., Tomii, K. & Katoh, K. Parallelization of MAFFT for large-scale multiple sequence alignments. Bioinformatics 34, 2490–2492 (2018).

38. Capella-Gutiérrez, S., Silla-Martínez, J.M. & Gabaldón, T. trimAl: a tool for automated alignment trimming in large-scale phylogenetic analyses. Bioinformatics 25, 1972–1973 (2009).

39. Stamatakis, A. RAxML version 8: a tool for phylogenetic analysis and post-analysis of large phylogenies. Bioinformatics 30, 1312–1313 (2014).

40. Yu, G., Smith, D.K., Zhu, H., Guan, Y. & Lam, T.T.-Y. ggtree: an r package for visualization and annotation of phylogenetic trees with their covariates and other associated data. Methods in Ecology and Evolution 8, 28–36 (2017).

41. Team, R. RStudio: Integrated Development Environment for R. RStudio, PBC (2020).

42. Team, R.C. R: A Language and Environment for Statistical Computing. R Foundation for Statistical Computing (2020).

43. Guo, X. et al. CNSA: a data repository for archiving omics data. Database 2020 (2020).

44. Chen, F.Z. et al. CNGBdb: China National GeneBank DataBase. Yi Chuan 42, 799–809 (2020).

